# Ericoid mycorrhizal diversity increases with soil age and progressive phosphorus limitation across a 4.1 million-year chronosequence

**DOI:** 10.1101/2020.08.27.270413

**Authors:** Devin R. Leopold, Kabir G. Peay, Peter M. Vitousek, Tadashi Fukami

## Abstract

Ericaceous plants rely on ericoid mycorrhizal fungi for nutrient acquisition. However, the factors that affect the composition and structure of these fungal communities remain largely unknown. Here, we use a 4.1-myr soil chronosequence in Hawaii to test the hypothesis that changes in nutrient availability with soil age determine the diversity and species composition of fungi associated with ericoid roots. We sampled roots of a native Hawaiian plant, *Vaccinium calycinum*, and used DNA metabarcoding to quantify changes in fungal diversity and species composition. We also used a fertilization experiment at the youngest and oldest sites to assess the importance of nutrient limitation. We found an increase in diversity and a clear pattern of species turnover across the chronosequence, driven largely by putative ericoid mycorrhizal fungi. Fertilization with nitrogen at the youngest site and phosphorus at the oldest site reduced total fungal diversity, suggesting a direct role of nutrient limitation. Our results also reveal the presence of novel fungal species associated with Hawaiian Ericaceae and suggest a greater importance of phosphorus availability for communities of ericoid mycorrhizal fungi than is generally assumed.

## Introduction

Most plants in the family Ericaceae host a diverse assemblage of root-associated fungi and form a unique mycorrhizal symbiosis known as an ericoid mycorrhiza (ErM; D.R. Leopold, 2016; Perotto et al., 2012). The ErM symbiosis allows ericaceous plants to proliferate in harsh environments where acidic soils, low temperatures, or excessive soil moisture slow the degradation of organic matter and limit mineral nutrient availability (Read 1991; Cairney and Meharg 2003; Mitchell and Gibson 2006). In these harsh habitats, which can be found on every continent except Antarctica (Kohout 2017), ericoid mycorrhizal fungi (ErMF) facilitate nutrient cycling by degrading complex organic matter and providing their hosts with access to nutrient pools that would be unavailable otherwise (Nasholm *et al*. 1998; Wurzburger, Higgins and Hendrick 2012; Adamczyk *et al*. 2016; Perotto, Daghino and Martino 2018). Ericaceous plants and ErMF also contribute to carbon sequestration in these environments through the production recalcitrant plant litter and hyphal necromass (Clemmensen *et al*. 2013, 2015). However, despite the ecological importance of the ErM symbiosis, and significant functional variation within and among ErMF species (Cairney *et al*. 2000; Whittaker and Cairney 2001; Grelet *et al*. 2009b; Wurzburger, Higgins and Hendrick 2012), little is known about how factors such as soil nutrient availability affect the community composition of ErMF and other fungi associated with ericoid roots.

Currently, most data on ErMF originates from temperate and boreal regions, where cooler temperatures and repeated glaciation throughout the Pleistocene have maintained widespread nitrogen (N) limitation, even in late-successional ecosystems (Tamm 1991; Vitousek and Howarth 1991). As a result, research on ErMF has focused on their ability to utilize various organic N sources and influence host N status (Leake and Read 1991; Nasholm *et al*. 1998; Xiao and Berch 1999; Cairney *et al*. 2000; Grelet *et al*. 2009b; Ishida and Nordin 2010). However, the distribution of ericaceous plants is not limited by these habitat conditions (Leopold 2016; Kohout 2017) and there is evidence that ErMF also facilitate the uptake of phosphorus (P) and other mineral nutrients for host plants (Pearson and Read 1973; Read 1983; Myers and Leake 1996). This suggests that the relative availability of limiting nutrients in soil, P as well as N, may influence ericaceous plant-fungal interactions and ErMF community composition (Hazard *et al*. 2014; Van Geel *et al*. 2020).

One source of natural variation in the availability of N and P in soil is pedogenesis, or long-term soil development. Over hundreds of thousands to millions of years, and in the absence of rejuvenating disturbance (Peltzer *et al*. 2010), pedogenesis causes predictable changes in ecosystem-level nutrient limitation, primarily through the accumulation of N from biological inputs and the progressive loss and occlusion of rock-derived nutrients, especially P (Walker and Syers 1976; Chadwick *et al*. 1999). These changes result in an initial, progressive phase, defined by N limitation of primary productivity and the rapid accumulation of organic matter; a mature phase, defined by maximal productivity and co-limitation by N and P; and a retrogressive phase, defined by P limitation and reduced rates of nutrient cycling (Vitousek and Farrington 1997; Richardson *et al*. 2004; Wardle 2004; Peltzer *et al*. 2010).

Because the direct observation of long-term soil development in a single site is not possible, soil chronosequences, or gradients of soil age in which other putative controlling factors (i.e., parent material, climate, vegetation type, etc.) are held reasonably constant, can provide insight into the effects of changes in nutrient availability with soil age (Fukami and Wardle 2005; Lambers *et al*. 2008; Walker *et al*. 2010; Dickie *et al*. 2013). Soil chronosequence studies of arbuscular mycorrhizal fungi (AMF) have suggested direct N and P limitation of fungi in young and old ecosystems, respectively (Treseder and Allen 2002), and a peak in AMF richness at fertile, middle-aged sites (Krüger *et al*. 2015). However, AMF are primarily involved in mineral nutrient uptake (Smith and Read 2006) and appear to lack the extracellular enzymes needed to access the organic nutrient pools available to some ErMF (Tisserant *et al*. 2013). ErMF diversity could respond differently to changing mineral nutrient availability with soil aging, but the relative importance of N and P limitation for ErMF is currently unknown. Furthermore, despite adaptation of the ErM symbiosis to soils with low mineral nutrient availability, studies of ErMF from retrogressive, P-limited habitats are lacking (Dickie *et al*. 2013).

To explore the effects of long-term soil development on ErMF, and the broader community of ericaceous root-associated fungi, we sampled roots of a single ericaceous plant species, *Vaccinium calycinum*, across a 4.1 myr soil chronosequence in the Hawaiian Islands, known as the Long Substrate Age Gradient (LSAG; Vitousek 2004). The LSAG is a useful system to study ErMF for three reasons. First, *V. calycinum*, a common host plant species, is present at all LSAG sites. The prevalence of a common host species across the chronosequence controls for changes in host identity, which can influence the composition of mycorrhizal symbiont communities (Martínez-García *et al*. 2015), though intraspecific variation in host traits affecting plant-fungal interactions are also possible (Johnson *et al*. 2010). Second, vegetation at LSAG sites is dominated by a single canopy tree species and a common suite of understory plants (Kitayama and Mueller-Dombois 1995), which all form mycorrhizal associations with AMF (Koske, Gemma and Flynn 1992). The only other ericaceous species present at any of the LSAG sites, *Vaccinium dentatum* and *Leptecophylla tameiameiae*, occur very sparsely at all sites, which further limits the potential impact of changes in host vegetation along the chronosequence. Third, changes in nutrient limitation across the LSAG have been experimentally demonstrated through N and P fertilizer addition experiments (Vitousek 2004). Because these fertilization experiments are long-term and ongoing at the oldest and youngest LSAG sites, this system presents a unique opportunity to experimentally test whether ecosystem-level nutrient limitation (i.e., N and P availability in young and old sites, respectively) plays a role in structuring communities of ericaceous root-associated fungi.

We began with the hypothesis that the diversity of fungi associated with ericaceous roots increases throughout long-term soil development, with ErMF diversity increasing owing to the accumulation of complex organic nutrient pools. We predicted that diversity would increase most rapidly during the early stages of pedogenesis due to the formation and development of an organic soil horizon and that an increase would continue into the retrogressive stages as soil weathering and occlusion of mineral nutrients promotes niche partitioning in the rhizosphere (Turner 2008; DeForest and Scott 2010). We also used long-term fertilization experiments to test the hypothesis that nutrient limitation of primary productivity, N at the youngest site and P at the oldest site, shapes the species composition of ErMF and other ericaceous root-associated fungi.

## Materials and Methods

### Study system

The LSAG chronosequence consists of six study sites that were established to address questions about the mechanisms that control nutrient cycling and limitation throughout long-term soil development by exploiting the increasing age of Hawaiian volcanoes with increasing distance from the active hot spot (Vitousek 2004). Samples for the study reported here were collected from five LSAG sites ranging in substrate age from 300 yr to 4.1 myr (Fig. 1). One site (1.4 myr) was not sampled due to logistical constraints. In this system, substrate age is an estimate of how long ago the parent substrate was deposited by volcanic activity. The locations of the LSAG sites were chosen to minimize variation in state factors other than the age of the soil substrate. All sites share similar initial parent substrate, occur on the constructional slope of a shield volcano, are *ca*. 1200 m asl, receive *ca*. 2500 mm rain annually, and have a mean annual temperature of *ca*. 16 °C (for detailed site descriptions, see Crews et al., 1995; Vitousek, 2004). Biotic variation is also highly constrained across the LSAG. Vegetation consists of native forest dominated by the canopy tree species *Meterosideros polymorpha*, with an understory that includes *Cheirodendron trigynum, Cibotium* spp. (tree ferns), *Coprosma* spp., *Ilex anomala, Myrsine* spp., and *V. calycinum* (Kitayama and Mueller-Dombois 1995).

**Figure 1:**
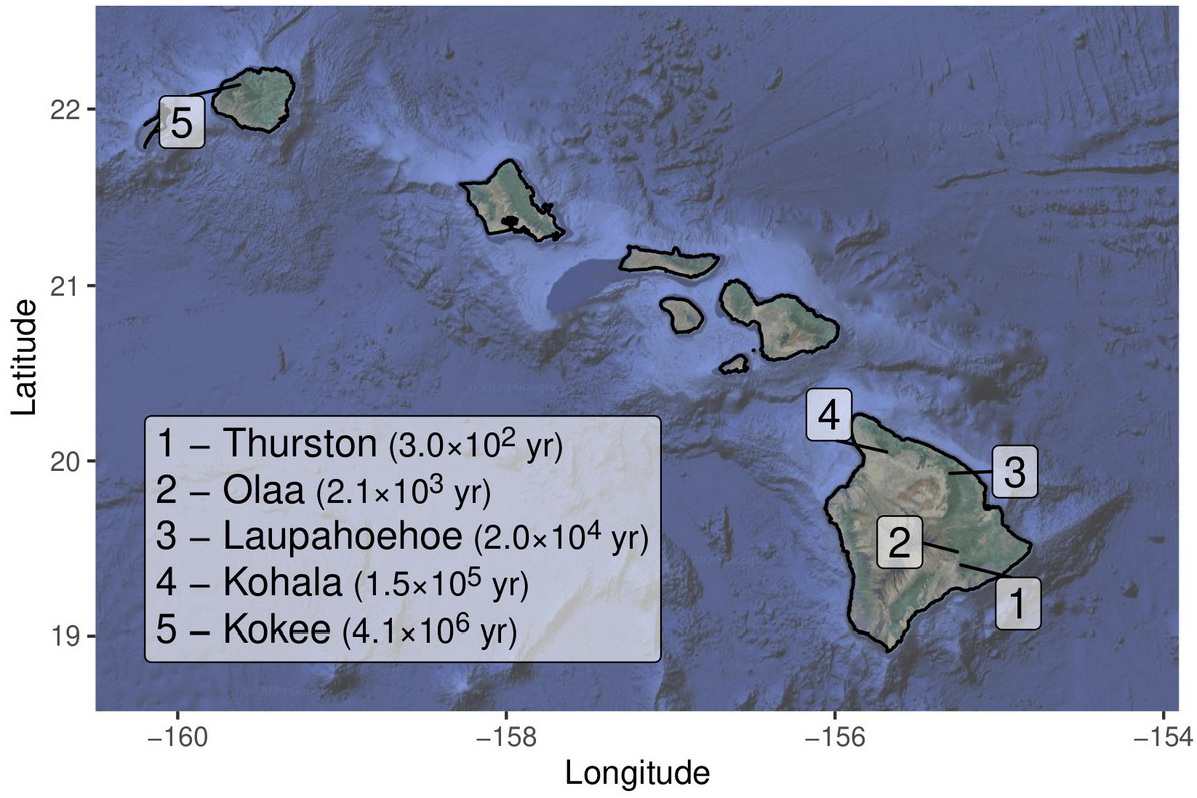
Map of the main Hawaiian Islands showing the locations and ages of the 5 sites in the Long Substrate Age Gradient chronosequence used in the current study.

In addition to the observational data collected across the LSAG chronosequence, fertilizer addition experiments were established in some LSAG sites to experimentally test hypotheses concerning the dynamics of nutrient limitation to ecosystem processes over geologic time scales. Fertilization plots sampled for the current study were established at the 300-yr site in 1985 (Vitousek *et al*. 1993) and at the 4.1-myr site in 1991 (Herbert and Fownes 1995), and so had been ongoing for 30 and 23 years, respectively, at the time of this sampling. At each site, 16 plots (15 × 15 m) were randomly assigned to either an unfertilized control or treatments that received fertilizer in the form of N (initially half ammonium nitrate and half urea, since 2002 just urea), P (triple superphosphate), or both N and P. Fertilizer was applied twice a year until 2002 and biannually thereafter at a consistent annual rate of 100 kg ha^−1^ yr^−1^.

### Sample collection

Sampling occurred in August 2014, with the exception of the nutrient addition plots at the 300 yr site, which were sampled in August 2013 as part of a preliminary feasibility study. To account for possible inter-annual variation, an additional 12 plants from non-fertilized plots at the 300 yr site were collected in 2014. Initial analyses suggested that samples from unfertilized plots collected at the 300 yr site in different years had similar fungal communities. Nonetheless, comparisons across the chronosequence reported here utilize only the 2014 samples and analyses of samples from fertilizer plots involved comparisons within each site, using the 2013 samples for the 300 yr site and the 2014 samples for the 4.1 myr site.

At each LSAG site, 12 mature *V. calycinum* plants were randomly selected within a 200 × 200 m area. In the fertilizer addition plots at the 300 yr and 4.1 myr sites, two plants were sampled in each 15 × 15 m plot, resulting in 32 samples per site (8 samples per fertilizer treatment in each site). For each plant, a portion of the root system and the adhering soil was removed with a hand trowel and bagged for transport. Samples were refrigerated (4 °C) until they could be processed, always within 48 hrs of collection. Fine terminal roots were manually separated from the soil and rinsed in tap water to remove all visible soil particles and 12 segments, *ca*. 2 cm each, were randomly selected and pooled for each plant. Pooled root samples were surface sterilized by sequential vortexing for 1 minute in sterile water, 70% EtOH, 50% household bleach, and then rinsed 3 times in sterile water. Surface sterilized roots were then stored at −80 °C.

### Fungal community metabarcoding

Frozen roots were homogenized by bead beating in CTAB lysis buffer and total DNA was extracted using the Nucleospin Plant II kit (Macherey-Nagel). In order to provide negative controls for Illumina sequencing, 4 sterile bead beating tubes were processed in parallel with the tubes containing root samples. Following DNA extraction, the first half of the internal transcribed spacer region (ITS1) of fungal nrDNA was amplified using Illumina fusion PCR primers. Primers included the ITS1 primers ITS1F (forward) or ITS2 (reverse), modified to include Illumina adapters and a sample specific 12 bp, error-correcting Golay barcode (Smith and Peay 2014). PCR was carried out in 25 μl reactions using 1 μl of template DNA (diluted 1:20), 0.5 μl of each 10 μM primer, 12.5 μl OneTaq Hot-Start 2X Master Mix (New England Biolabs) and a cycling program consisting of: initial denaturing at 94 °C (1 min), 30 cycles of 95 °C (30 sec), 52 °C (30 sec) and 68 °C (30 sec) and a final elongation stage of 68 °C (5 min). PCR reactions were carried out in triplicate for each sample and were individually checked for successful amplification using gel electrophoresis. The individual PCR reactions for each sample were cleaned and normalized to 2.5 ng μl^−1^ using a Just-a-Plate, 96 well normalization and purification plate (Charm Biotech). Samples were then pooled and sequenced using Illumina MiSeq paired-end sequencing (2 x 300 bp) at the Stanford Functional Genomics Facility. Raw sequence data were deposited in the NCBI short read archive under BioProject ID PRJNA548137.

### Bioinformatics

Preliminary analyses suggested that the reverse read quality was poor, so only the forward reads were used for all analyses. Raw reads were first trimmed to remove 3’ gene primer and sequencing adapter contamination using cutadapt (Martin 2011). The trimmed reads were then pooled and denoised using *DADA2* (Callahan *et al*. 2016), excluding reads with an expected error rate > 2 bp, a length < 75 bp, and putative chimeric sequences. Denoised reads were then collapsed to 99% operational taxonomic units (OTUs) using agglomerative, single-linkage clustering implemented in *DECIPHER* (Wright, Erik 2016). The OTU table was then filtered of likely contaminants by removing any OTU with an average abundance across all samples that was less than the maximum abundance in any of the negative controls.

Taxonomic predictions for OTUs were made using the *DADA2* implementation of the RDP classifier (Wang *et al*. 2007) trained on the UNITE v8.2 species hypothesis database (Abarenkov, Kessy; Zirk, Allan; Piirmann, Timo; Pöhönen, Raivo; Ivanov, Filipp; Nilsson, R. Henrik; Kõljalg 2020). Because a prominent group of ErMF, known as the *Rhizoscyphus ericae* aggregate, or REA (≡ *Hymenoscyphus ericae* aggregate; Vrålstad, Fossheim and Schumacher 2000; Vrålstad, Myhre and Schumacher 2002; Hambleton and Sigler 2005), has undergone numerous taxonomic revisions, we manually curated the relevant taxonomy in the training data. We resolved inconsistencies in this clade following the most recent updates reported by Fehrer et al. (2019), which consolidated members of the REA in the genus *Hyaloscypha*.

Identification of potential ErMF in the OTU database was complicated by two factors. First, the short read length and high variability of the ITS region, combined with limited representation of both Hawaiian root / soil associated fungi and fungi associated with ericaceous plants in current reference databases resulted in low confidence for fine scale taxonomic assignments. Second, there is significant uncertainty about the taxonomic range of fungal species capable of forming ErM. We chose an inclusive approach, defining the subset of putative ErMF to include all OTUs assigned to fungal orders reported to include ericoid fungi (Leopold 2016). This approach is likely to include many non-mycorrhizal endophytes, particularly in the order Helotiales, however some of these species likely have overlapping functional attributes with ErMF (Newsham 2011) and it is not currently possible to distinguish ErMF taxa with sequence data alone. An alternative approach, manually curating putative ErMF OTUs by looking at the sources of the individual sequences comprising the closest matching UNITE species hypotheses yielded qualitatively similar results. We focus here on the approach using fungal orders as it is more easily replicated.

### Statistical analysis of chronosequence data

To assess if fungal diversity changed across the LSAG, we estimated alpha diversity at the sample level for both the complete data set and the subset of putative ErMF. To account for unequal sequencing depth among samples we estimated alpha diversity using Hill numbers, or the effective number of species, at a sampling depth of 5000 sequences per sample using the R-package *iNEXT* (Chao *et al*. 2014; Hsieh, Ma and Chao 2016). We calculated Hill numbers for each sample using a scaling of 0 (equivalent to species richness) and a scaling of 1 (equivalent to the exponent of Shannon entropy, or Shannon diversity) and analyzed the results in parallel. Variation in alpha diversity among chronosequence sites was tested using ANOVAs and *F*-tests, followed by Tukey’s HSD post-hoc tests.

To determine how the composition of fungal communities changed across the LSAG, we first calculated Bray-Curtis dissimilarity among samples using the proportional abundance of OTUs in each sample to account for unequal sampling depth (McMurdie and Holmes 2014). We initially tested alternative dissimilarity metrics, including the binary Jaccard distance, the probabilistic Raup-Crick distance, and the information-theoretic based Jensen-Shannon Divergence, and found that all produced qualitatively similar results to those obtained with Bray-Curtis dissimilarity, which we will focus on here. Dissimilarities were visualized using nonmetric multidimensional scaling (NMDS) implemented in the R-package *vegan* (Oksanen *et al*. 2017). Contours representing increasing site age were fit to the 2-dimensional NMDS ordination, using the vegan function ordisurf, to aid interpretation of species turnover across the chronosequence. The significance of changes in community composition were tested with a permutational multivariate analysis of variance (Anderson 2001). Both log10-transformed site age and site identity as a factor were used as predictors in perMANOVAs to assess both the significance of variation in community composition among sites and the proportion of this variation that could be explained by turnover along the chronosequence. OTUs occurring at individual LSAG sites with greater abundance and more often than expected by chance alone were identified using indicator species analysis (Dufrêne and Legendre 1997) using the R-package *labdsv* (Roberts 2013), and visualized on a bipartite graph of the 50 most abundant OTUs.

### Statistical analysis of fertilizer plot data

To test the effects of fertilization on fungal composition and diversity, we analyzed both the complete data set and the subset of putative ErMF OTUs from the fertilizer addition plots at the 300 yr and 4.1 myr LSAG sites. We followed the procedure described above to estimate fungal richness and Shannon diversity and then used ANOVAs and *F*-tests to assess the direct and interactive effects of N and P addition. Preliminary analysis indicated that the fungal richness and diversity associated with pairs of plants sampled from the same plot were not more similar than those between plots with the same fertilizer treatments, so analyses were conducted at the level of individual plants (n = 32), rather than fertilization plots (n = 16). To test whether community composition was affected by fertilization, we used NMDS and permutational MANOVAs of Bray-Curtis dissimilarity calculated on proportional OTU abundance.

### Reproducibility of analyses

All statistical analyses were conducted in the R v4.2 statistical computing environment (R Core Team 2020), using the package *tidyverse* (Wickham *et al*. 2019) for general data manipulation and *phyloseq* (McMurdie and Holmes 2013) to manipulate OTU tables and associated metadata. Code for reproducing all bioinformatic processing, statistical analyses and figures, along with all necessary sample metadata, has been publicly archived (DOI: 10.5281/zenodo.3979769).

## Results

### Identification of fungal OTUs

From 4.6 M raw reads, we retained 2.2 M high-quality sequences after bioinformatic processing, from which we identified 685 fungal OTUs. The subset of putative ErMF included 30.8% of the total OTUs, and accounted for 66.5% of all sequences. Many taxa could not be confidently assigned to genera or family (Fig. 2), and the lack of high similarity matches for many OTUs in the UNITE database suggests the presence of many novel taxa. For example, the most abundant taxon we identified (OTU.1) shares only 85% sequence similarity with the nearest sequence in the UNITE database and could not be confidently assigned beyond the order Helotiales. The second most abundant taxon (OTU.2) was assigned to the order Trechisporales and shares 99% sequence similarity with a UNITE species hypothesis that contains only a few individuals sequenced without culturing from the roots of plants in the Orchidaceae and Clustaceae, originating from Eastern Africa and Taiwan. However, the next closest UNITE species hypothesis in the Trechisporales shares only 75% sequence similarity, suggesting that this is could be a specious taxonomic assignment (see discussion, below). We did identify numerous taxa with affinities to the globally widespread *Rhizoscyphus ericae* aggregate (≡ *Hymenoscyphus ericae* aggregate; Vrålstad, Fossheim and Schumacher 2000; Vrålstad, Myhre and Schumacher 2002; Hambleton and Sigler 2005), including the common ErMF, *Hyaloscypha hepaticicola* (≡ *Rhizoscyphus ericae* & *≡Pezoloma ericae;* Fehrer et al. 2019), though this taxon was notably absent at the youngest site (Fig. 2). The common ErMF species, *Oidiodendron maius*, was only observed in very low abundance in the sequence data, primarily in samples from the oldest chronosequence site. No members of the Sebacinales were observed, despite their common occurrence as root endophytes and ErMF on ericaceous plants (Selosse *et al*. 2007; Weiß *et al*. 2011).

**Figure 2:**
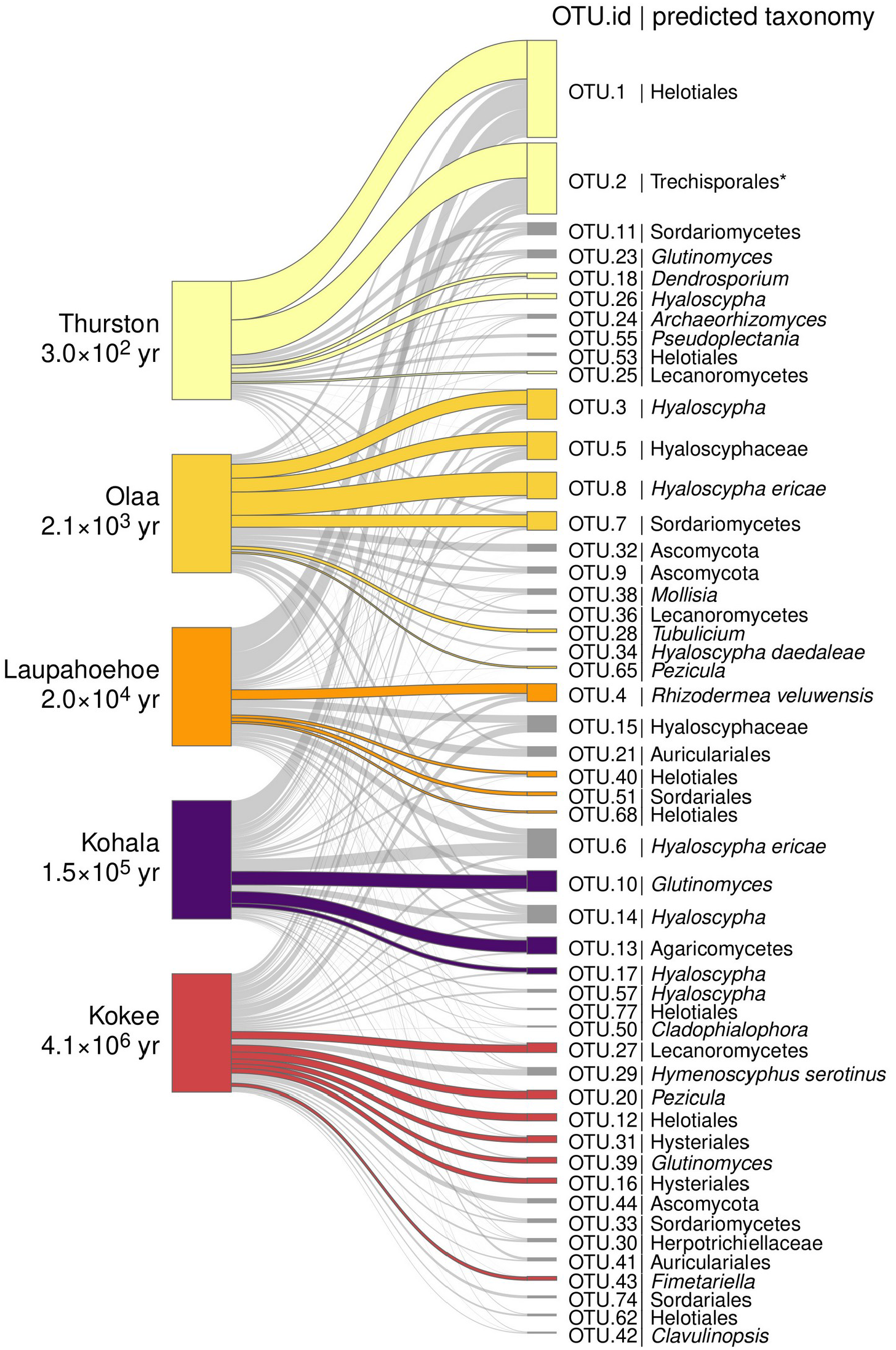
Bipartite network showing the relative abundances of the 50 most common fungal OTUs associated with the roots of *Vaccinium calycinum* across the Long Substrate Age Gradient. The size of nodes representing OTUs (right) is proportional to their mean abundance in the full data set and the width of connections between OTUs and sites (left) is proportional to their relative abundance at each site. Colored OTU nodes and connections identify significant site associations (indicator species analysis). Taxonomic predictions for each OTU are indicated following the unique OTU identifier.

### Site age effects on fungal diversity and composition

Fungal richness and diversity were greatest in *V. calycinum* roots collected from the oldest chronosequence sites (Fig. 3). When all fungi were considered together (Fig. 3a), variation in OTU richness (*F*_4,55_ = 5.14; *p* = 0.001) was primarily the result of lower richness in the more fertile, middle-aged sites, while variation Shannon diversity (*F*_4,55_ = 11.9; *p* < 0.001) was primarily due to greater richness at the oldest site. For putative ErMF (Fig. 3b), variation in OTU richness (*F*_4,54_ = 5.62; *p* < 0.001) was the result of an increase at the two oldest sites, relative to the younger sites, while variation in Shannon diversity (*F*_4,54_ = 11.4; *p* < 0.001) was the result of a progressive increase with site age.

**Figure 3:**
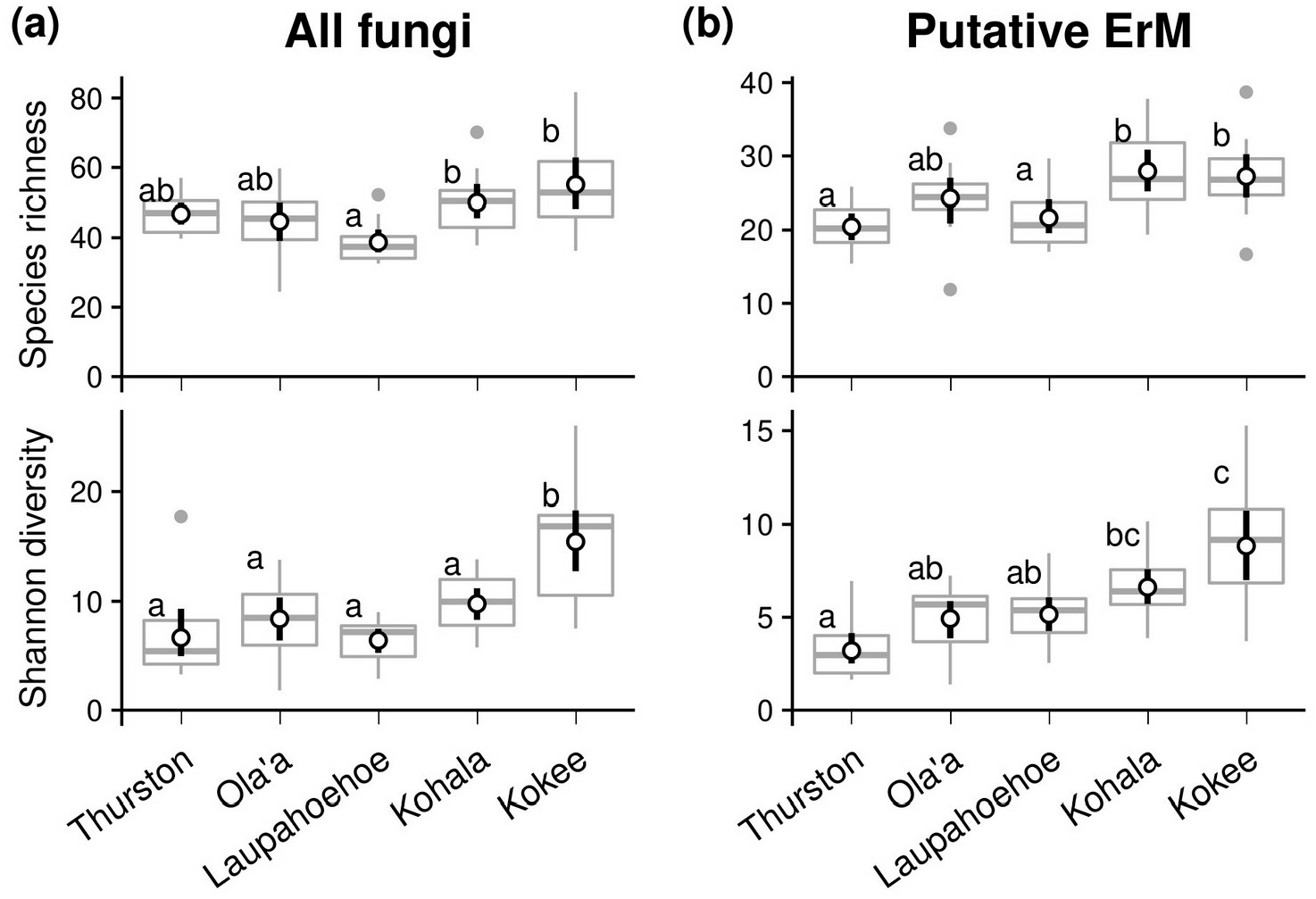
Variation in species richness and Shannon diversity of (a) all fungi and (b) putative ErMF associated with *Vaccinium calycinum* roots across the Long Substrate Age Gradient. Boxplots show the distribution of the point estimates of richness and diversity for 12 replicate samples at each site. Points indicate the predicted mean (± 95% bootstrapped CI) species richness or diversity. Letters indicate significant pairwise differences in post-hoc comparisons.

Fungal communities associated with *V. calycinum* roots differed in composition among chronosequence sites for all fungi (*F*_4,55_ = 6.92, *r^2^* = 0.33, *p* < 0.001) and the subset of putative ErMF (*F*_4,55_ = 9.42, *r*^2^ = 0.40, *p* < 0.001). Similarly aged sites tended to have more similar OTU compositions (Fig. 4), and site age explained roughly a third of the variation in OTU composition that was attributed to variation among sites, both for all fungi (*F*_1,58_ = 7.64, *r*^2^ = 0.12, *p* < 0.001) and putative ErMF (*F*_1,58_ = 9.41 *r*^2^ = 0.14, *p* < 0.001). In addition, while many of the most abundant OTUs were broadly distributed across the chronosequence, occurring in most or all sites, distribution patterns varied among taxa and many could be identified as being significantly associated with individual LSAG sites (Fig. 2).

**Figure 4:**
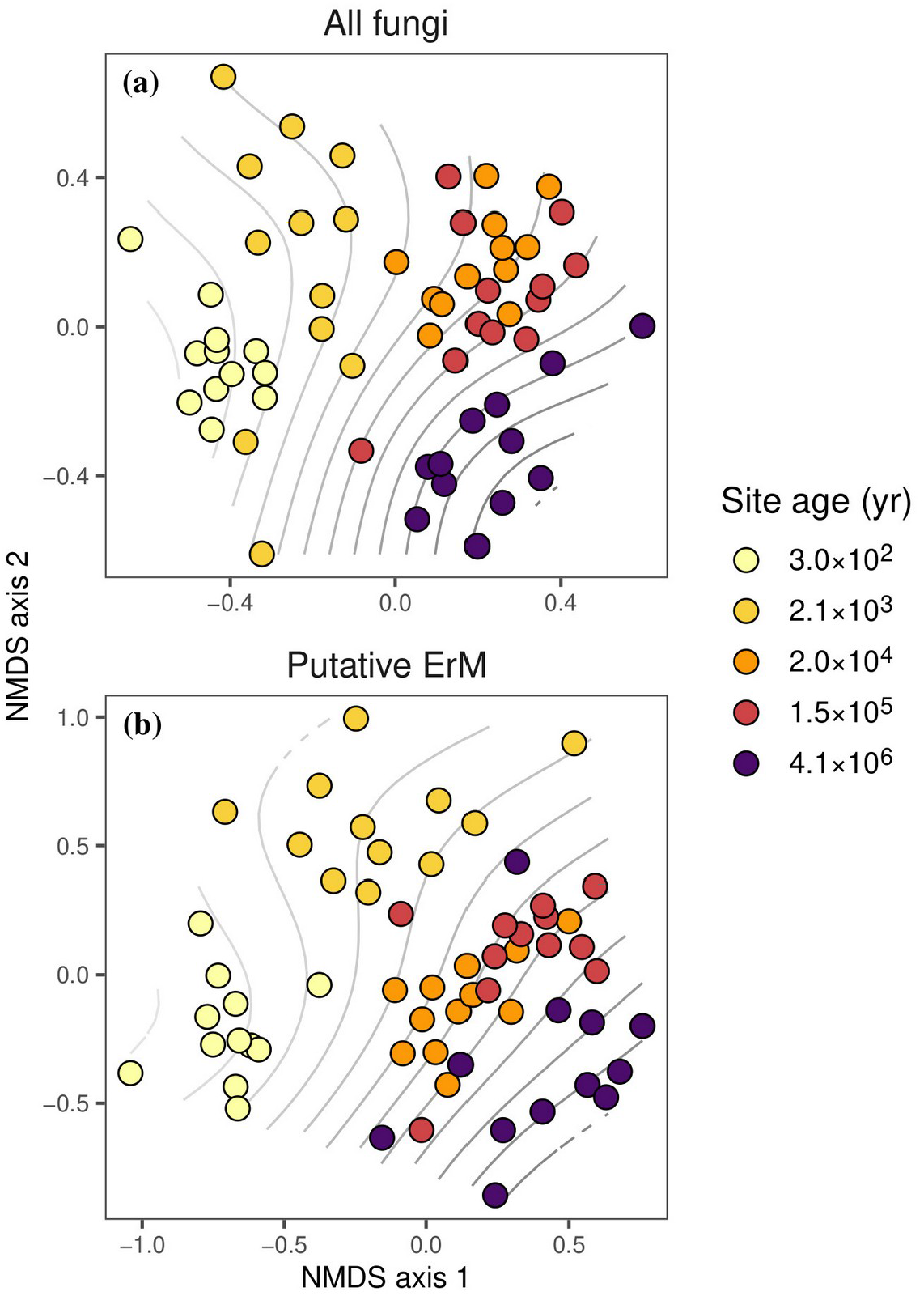
Non-metric multidimensional scaling of community dissimilarity (Jensen Shannon distance) for (a) all fungi (stress = 0.26) and (b) putative ErMF (stress=0.25) associated with *Vaccinium calycinum* roots collected across a 4.1-myr chronosequence. Points represent individual plants, with color indicating the age of the site where the sample was collected. Contours indicate a smooth surface of log_10_-transformed site age fit to the ordination space, where increasing site age is indicated by darker contours.

### Fertilization effects on fungal composition and diversity

The effect of fertilization on fungal richness and diversity depended on site (Fig. 5; Tables S1 & S2). At the 300 yr-old site, the addition of N reduced both richness and Shannon diversity for all fungi (*F*_1,28_ = 10.1; *p* = 0.004 & *F*_1,28_ = 5.95; *p* = 0.02, respectively). For putative ErMF, the addition of N at the 300 yr-old site also reduced OTU richness (*F*_1,28_ = 12.9; *p* = 0.001), and marginally reduced Shannon diversity (*F*_1,28_ = 3.37; *p* = 0.08). In contrast, at the 4.1 myr site it was the addition of P that reduced OTU richness and Shannon diversity for all fungi (*F*_1,28_ = 6.65; *p* = 0.02 & *F*_1,28_ = 15.8; *p* < 0.001, respectively) and putative ErMF (*F*_1,28_ = 3.60; *p* = 0.07 & *F*_1,28_ = 8.7; *p* = 0.006, respectively).

**Figure 5:**
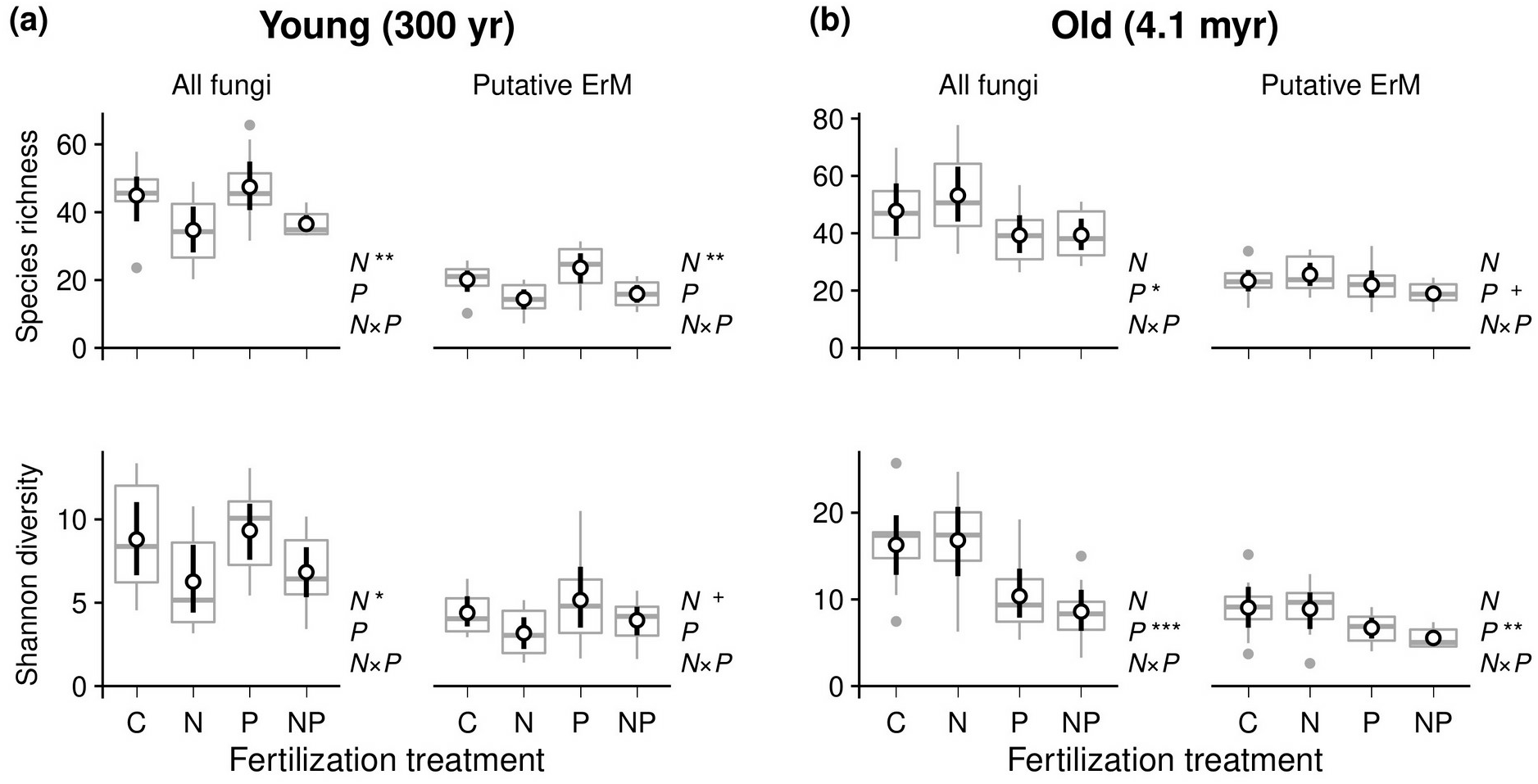
The effect of fertilization with nitrogen (N), phosphorus (P), or both (NP) on the species richness and Shannon diversity of fungal OTUs associated with the roots of *Vaccinium calycinum* at (a) the 300 yr and (b) the 4.1 myr (b & d) sites in the Long Substrate Age Gradient chronosequence. Boxplots show the distribution of individual replicate samples and points indicate the predicted mean (± 95% CI) species richness or diversity for each treatment of the unfertilized control (C). Significant direct and interactive effects of N and P additions are indicated to the right of each panel, where + *p* < 0.1, * *p* < 0.05, ** *p* < 0.0.1, and *** *p* < 0.0.1.

In contrast to richness and diversity, fungal community composition was not primarily impacted by addition of the limiting nutrient at each site (Table 1; Fig. S1). Instead, both N and P additions affected fungal composition, but P addition had the greatest impact on composition at both sites, for all fungi and the subset of putative ErMF.

**Table 1:** Results of permutational MANOVAs examining the effect of fertilizer addition treatments on the composition of (a) all fungi or (b) putative ErMF associated with *V. calycinum* at the 300 yr and 4.1 myr sites in the Long Substrate Age Gradient chronosequence.

**a) All fungi**
| **Site** | **Treatment** | ***F*** | ***r*<sup>2</sup>** | ***P*** |
| --- | --- | --- | --- | --- |
| Young (300 yr) | Full model | 1.876 | 0.167 | < 0.001 |
|  | N | 2.284 | 0.068 | 0.003 |
|  | P | 2.717 | 0.081 | 0.001 |
|  | N:P | 0.651 | 0.019 | 0.913 |
| Old (4.1 myr) | Full model | 1.767 | 0.159 | < 0.001 |
|  | N | 1.588 | 0.047 | 0.020 |
|  | P | 2.647 | 0.080 | < 0.001 |
|  | N:P | 1.066 | 0.032 | 0.345 |

**b) Putative ErMF**
| **Site** | **Treatment** | ***F*** | ***r*<sup>2</sup>** | ***P*** |
| --- | --- | --- | --- | --- |
| Young (300 yr) | Full model | 1.787 | 0.160 | 0.007 |
|  | N | 1.488 | 0.044 | 0.130 |
|  | P | 3.283 | 0.099 | < 0.001 |
|  | N:P | 0.549 | 0.016 | 0.891 |
| Old (4.1 myr) | Full model | 2.324 | 0.199 | < 0.001 |
|  | N | 1.782 | 0.051 | 0.028 |
|  | P | 3.592 | 0.103 | < 0.001 |
|  | N:P | 1.600 | 0.046 | 0.060 |

## Discussion

Consistent with our initial hypothesis, we found that *V. calycinum* associates with more putative ErMF in sites with older soils. This pattern was not simply the result of increasing richness, but also greater evenness, which could reflect increased niche partitioning in response to the accumulation of soil organic matter and the development of a structured organic soil horizon during pedogenesis (Goh *et al*. 1976; Torn *et al*. 1997). Although speculative, this explanation is consistent with the fact that ErMF taxa can vary in their affinity for specific nutrient sources (Cairney *et al*. 2000; Whittaker and Cairney 2001; Grelet, Meharg and Alexander 2005; Grelet *et al*. 2009b) and can be spatially structured across soil layers (Wurzburger, Higgins and Hendrick 2012). However, it is also possible that dispersal limitation (Cline and Zak 2014) and *in-situ* diversification (Gillespie 2016) contributed to the greater diversity observed in older sites and manipulative experiments would be needed to quantify the role of niche partitioning relative to other factors.

Results from our fertilization experiment suggest that soil-age related nutrient limitation influenced fungal richness and diversity. Specifically, we found that richness and diversity decreased in response to the addition of N and P at the youngest and oldest chronosequence sites, respectively. Previous studies have shown that plant growth is limited by N at the 300 yr site and P at the 4.1 myr site (Vitousek 2004), and P limits rates of litter decomposition and soil organic matter processing in the 4.1 myr site (Hobbie and Vitousek 2000; Reed, Vitousek and Cleveland 2011). Moreover, Treseder and Allen (2002) found that growth of arbuscular mycorrhizal fungi associating with non-ericaceous plants in this system is directly N-limited at the 300 yr site and P-limited at the 4.1 myr site. Our results could suggest that ErMF, in contrast with arbuscular mycorrhizal fungi, are not directly limited by N and P availability in the LSAG, possibly due to their ability to access recalcitrant organic nutrient pools (Read 1983, 1991; Perotto, Daghino and Martino 2018). Instead, fertilizer addition may cause the host plant to reallocate resources (i.e., carbon-rich photosynthates) away from root-associated symbionts (Olsson, Rahm and Aliasgharzad 2010; Kiers *et al*. 2011; Konvalinková *et al*. 2017), resulting in reduced fungal richness and diversity in the rhizosphere. This possibility could have significant implications for stability of ERM symbioses in the context of anthropogenic nutrient deposition in the nutrient-poor habitats where ericaceous plant often thrive (Van Geel *et al*. 2020).

Although we observed a clear pattern of turnover in species composition across the soil chronosequence (Fig. 4), results from our fertilization experiment suggest that host and ecosystem-level nutrient limitation are not the primary determinants of fungal species composition in this system (Table 1). This could be because dispersal limitation restricts the pools of ErMF species present at each site (Hutton *et al*. 1997), limiting the effect of localized nutrient additions. Nonetheless, both N and P additions did influence fungal community composition at both sites, though the effects of P additions were consistently larger, even at the N-limited site. Previous work in this system has shown that fertilization with P at the 4.1 myr site significantly affects litter chemistry, whereas fertilization with N at the 300 yr site does not (Harrington, Fownes and Vitousek 2001). Because ericaceous root systems occur predominantly in the upper, organic soil layers, where ErMF directly access recalcitrant litter-nutrient inputs (Read, Leake and Perez-Moreno 2004), changes in litter chemistry may explain why the strongest fertilization effects we observed for putative ErMF with P addition at the 4.1 myr site. These results are in contrast with the common characterization of ErM symbioses as being primarily N driven (Read 1991) and highlight the relative lack of data from P-limited tropical systems.

We do not have the histological data to determine the morphology of fungal interactions with *V. calycinum* and definitively determine the mycorrhizal status of the fungal OTUs we identified. However, our results suggest that many novel ErMF species could occur in this system. For example, the two most abundant OTUs in our study, which accounted for 32% of all reads across the chronosequence and 64% of the reads at the youngest site (Fig. 2), were not closely related to any well-described ErMF species. Although the most abundant OTU shared only 85% sequence similarity with the nearest taxon in the UNITE database, which comprises a large group of poorly classified Helotiales, many of the most similar taxa were isolated from the roots of ErM or ecto-mycorrhizal plants, which can host ErMF as endophytes or mycorrhizal fungi capable of forming both ecto- and ericoid mycorrhizae (Villarreal-Ruiz, Anderson and Alexander 2004; Vrålstad 2004; Grelet *et al*. 2009a; Villarreal-Ruiz *et al*. 2012; Vohník *et al*. 2013). There also is considerable reason to doubt the taxonomic prediction for the second most abundant OTU as Trechisporales. A mycorrhizal symbiont forming a unique, sheathed-ericoid mycorrhizal symbiosis in Norway was also originally thought to be most closely related to the Trechisporales (Vohník *et al*. 2012). This taxon was later identified as *Kurtia argillacea*, in the Hymenochaetales, based on multi-gene phylogenetic analyses, which showed that proper taxonomic placement using nrDNA genes was not possible for this taxon (Kolařík and Vohník 2018). Although taxon (OTU.2) does not share any substantive sequence similarity with the one available ITS sequence for *Kurtia argillacea*, it does share some characteristics, including an elevated GC content (57%), which Kolařík and Vohník (2018) suggest could be the result of a non-standard substitution rate, possibly related to the transition to a mycorrhizal lifestyle. Culture-based investigations of this taxon, and other novel taxa associated with Hawaiian ericaceous plant, are needed to definitively address the possibility that our data set includes novel ErMF. Although it is also possible that some of these species are non-mycorrhizal endophytes, this does not preclude a role in nutrient uptake.

In summary, we have shown that the diversity of fungi associated with the roots of ericaceous plants increases throughout long-term soil development largely due to increasing richness and evenness of putative ErMF taxa. Our results contrast with recent reports of a hump-shaped diversity pattern with increasing soil age for arbuscular mycorrhizal fungi (Krüger *et al*. 2015), suggesting that functional differences among types of mycorrhizal fungi may determine the response of root-associated fungi to long-term soil development. We have also shown that P availability can affect the species composition and community structure of ericoid fungi, particularly in older P-limited ecosystems. Our results suggest that further study of ErMF from P-limited systems or across long-term soil chronosequences may prove valuable for identifying the extent of taxonomic and functional diversity for this unique group of mycorrhizal fungi.

## Funding

This work as supported by Stanford University, Department of Biology, and student research grants from the American Society of Naturalists and the Mycological Society of America.

## Acknowledgments

We thank Michael Shintaku at the University of Hawaii, Hilo, College of Agriculture, Forestry and Natural Resource Management and Anne Veillet at the Evolutionary Genomics Core Facility for access to laboratory space and equipment. We also thank Matt Knope for feedback on earlier drafts of this manuscript. Site access was granted by the Hawaii Division of Land and Natural Resources, the Hawaii Experimental Tropical Forest, and Parker Ranch.

## Notes

### Competing Interest Statement

The authors have declared no competing interest.

