## Supplemental materials for "Ericoid mycorrhizal diversity increases with soil age and progressive phosphorus limitation across a 4.1 million-year chronosequence"

**Table S1:** Full ANOVA model results presented in Figure 5a in the main text, testing the effect of fertilization with nitrogen (N) or phosphorus (P) on the observed richness (a) and Shannon diversity (b) all fungi and the subset of putative ErMF at the 300 yr chronosequence site.

**a) Species richness**

| **Site** | **Treatment** | ***df*** | **Sum Sq** | **Mean Sq** | ***F*** | ***P*** |
|---|---|---|---|---|---|---|
| All fungi | N | 1 | 897.8 | 897.8 | 10.12 | 0.004 ** |
| | P | 1 | 36.35 | 36.35 | 0.410 | 0.527 |
| | N:P | 1 | 0.956 | 0.956 | 0.011 | 0.918 |
| | Residuals | 28 | 2,485 | 88.75 | | |
| ErMF | N | 1 | 358.2 | 358.2 | 12.95 | 0.001 *** |
| | P | 1 | 51.56 | 51.56 | 1.864 | 0.183 |
| | N:P | 1 | 8.235 | 8.235 | 0.298 | 0.590 |
| | Residuals | 28 | 774.7 | 27.67 | | |

**b) Shannon diversity**

| **Site** | **Treatment** | ***df*** | **Sum Sq** | **Mean Sq** | ***F*** | ***P*** |
|---|---|---|---|---|---|---|
| All fungi | N | 1 | 50.91 | 50.91 | 5.952 | 0.021 * |
| | P | 1 | 2.459 | 2.459 | 0.288 | 0.596 |
| | N:P | 1 | 0.000 | 0.000 | 0.000 | 0.999 |
| | Residuals | 28 | 239.5 | 8.552 | | |
| ErMF | N | 1 | 11.69 | 11.69 | 3.365 | 0.077 + |
| | P | 1 | 4.551 | 4.551 | 1.310 | 0.262 |
| | N:P | 1 | 0.000 | 0.000 | 0.000 | 0.997 |
| | Residuals | 28 | 97.25 | 3.473 | | |

**Table S2:** Full ANOVA model results presented in Figure 5b in the main text, testing the effect of fertilization with nitrogen (N) or phosphorus (P) on the observed richness (a) and Shannon diversity (b) all fungi and the subset of putative ErMF at the 4.1 myr chronosequence site.

**a) Species richness**

| Site | Treatment | *df* | Sum Sq | Mean Sq | *F* | *P* |
|---|---|---|---|---|---|---|
| All fungi | N | 1 | 61.37 | 61.37 | 0.410 | 0.527 |
| | P | 1 | 996.3 | 996.3 | 6.650 | 0.015 * |
| | N:P | 1 | 57.29 | 57.2 | 0.382 | 0.541 |
| | Residuals | 28 | 4195 | 149.8 | | |
| ErMF | N | 1 | 1.772 | 1.772 | 0.049 | 0.826 |
| | P | 1 | 129.5 | 129.5 | 3.595 | 0.068 + |
| | N:P | 1 | 54.94 | 54.94 | 1.525 | 0.227 |
| | Residuals | 28 | 1009 | 36.03 | | |

**b) Shannon diversity**

| Site | Treatment | *df* | Sum Sq | Mean Sq | *F* | *P* |
|---|---|---|---|---|---|---|
| All fungi | N | 1 | 3.327 | 3.327 | 0.131 | 0.720 |
| | P | 1 | 401.3 | 401.3 | 15.84 | 0.001 *** |
| | N:P | 1 | 10.78 | 10.78 | 0.426 | 0.519 |
| | Residuals | 28 | 709.5 | 25.34 | | |
| ErMF | N | 1 | 3.595 | 3.595 | 0.501 | 0.485 |
| | P | 1 | 64.34 | 64.34 | 8.965 | 0.006 ** |
| | N:P | 1 | 1.908 | 1.908 | 0.266 | 0.610 |
| | Residuals | 28 | 200.9 | 7.177 | | |

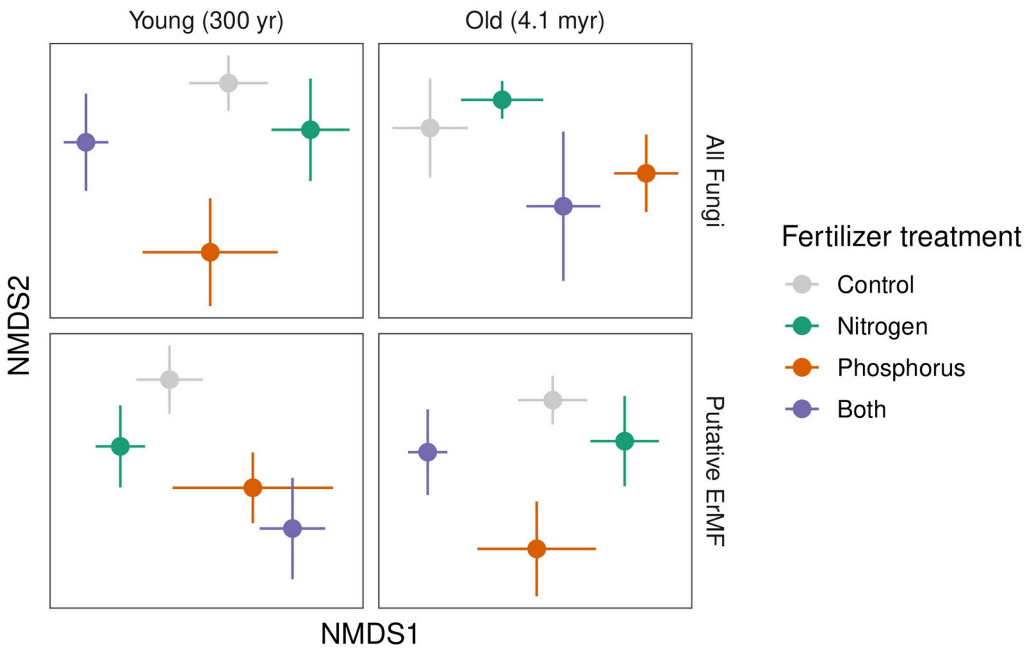


**Figure S1:** Non-metric multidimensional scaling of fungal community dissimilarity (Bray-Curtis) in long-term fertilizer addition plots at the N-limited, young site and the P-limited, old site. Colors indicate fertilizer addition treatments and points indicate centroids (± se) for each treatment.
